# Bidirectional connectivity changes in the frontotemporal areas reflect figure–ground reversal in multivoiced music

**DOI:** 10.1101/2024.05.08.593264

**Authors:** Chan Hee Kim, Jeong-Eun Seo, Chun Kee Chung

## Abstract

When listening to multivoiced music, it is possible to perceive distinct voices in differing pitch ranges independently. The melody that one focuses on becomes the figure, and other voices form the background. Shifting focus from one voice to another causes a figure–ground reversal, which may easily occur as a result of anticipation within repeated passages in a musical structure. We previously found that frontotemporal connectivity reflects changes in the melody of “Twinkle, Twinkle, Little Star” (TTLS) in Mozart’s 12 Variations K. 265. We hypothesized that if frontotemporal connectivity remains unchanged in repeated passages, the melody of TTLS remains the figure; however, if connectivity changes, the melody shifts toward the background. Our findings show that frontotemporal connectivity only changed during the final, fourth repetition. This was accompanied by a bidirectional correlation in frontotemporal regions, implying both top-down and bottom-up processes. We captured a momentary figure–ground reversal in the multivoiced texture of continuously changing and developing music.

## Introduction

The concepts of figure and ground in Gestalt psychology^1,2^ find parallels in the perceptually prominent and less prominent voices in multivoiced music. The neural processes underlying the perception of prominently heard voices (figure) and less prominently heard voices (ground), particularly when this prominence is naturally reversed while listening to music, remain inadequately investigated. Similar to visual stimuli (Fig. 1A), where attention can shift freely between a white star and a black ground, attention can shift to different voices within music. The auditory stream distinguishes the melodic line of the upper voice through the higher pitches^3,4^, but the lower voice can also be perceived as a figure, depending on its prominence in the musical context^5^. The melodic line of the highest voice tends to stand out perceptually due to its pitch range. However, the perceptual dominance of the melodic line is not fixed and can be influenced by listener’s attention and choice (Fig. 1B). The listeners may feel that the appearance of individual phrases change in relation to their focus on the voices^6^, in a type of figure–ground reversal between voices. The figure–ground relationship for upper and lower voices can be affected by the repetition of passages^7^. For later repetitions, the upper voice may be recognized as the ground and the lower voice as the figure (Fig. 1C). Musical structure, encompassing elements such as pitch, tonality, and harmonic structure, is implicitly learned through musical experience^8^. Recognizing similarities in musical structure helps developing hypotheses, categorizing inputs, and processing variables^9^. For example, while listing to the song “Twinkle, Twinkle, Little Star (TTLS)” of the ternary form, A (a + a) + B (b + a’) + B (b + a’), the following passages can easily be predicted (Fig. 2A, S1, and S2).

**Fig. 1.**
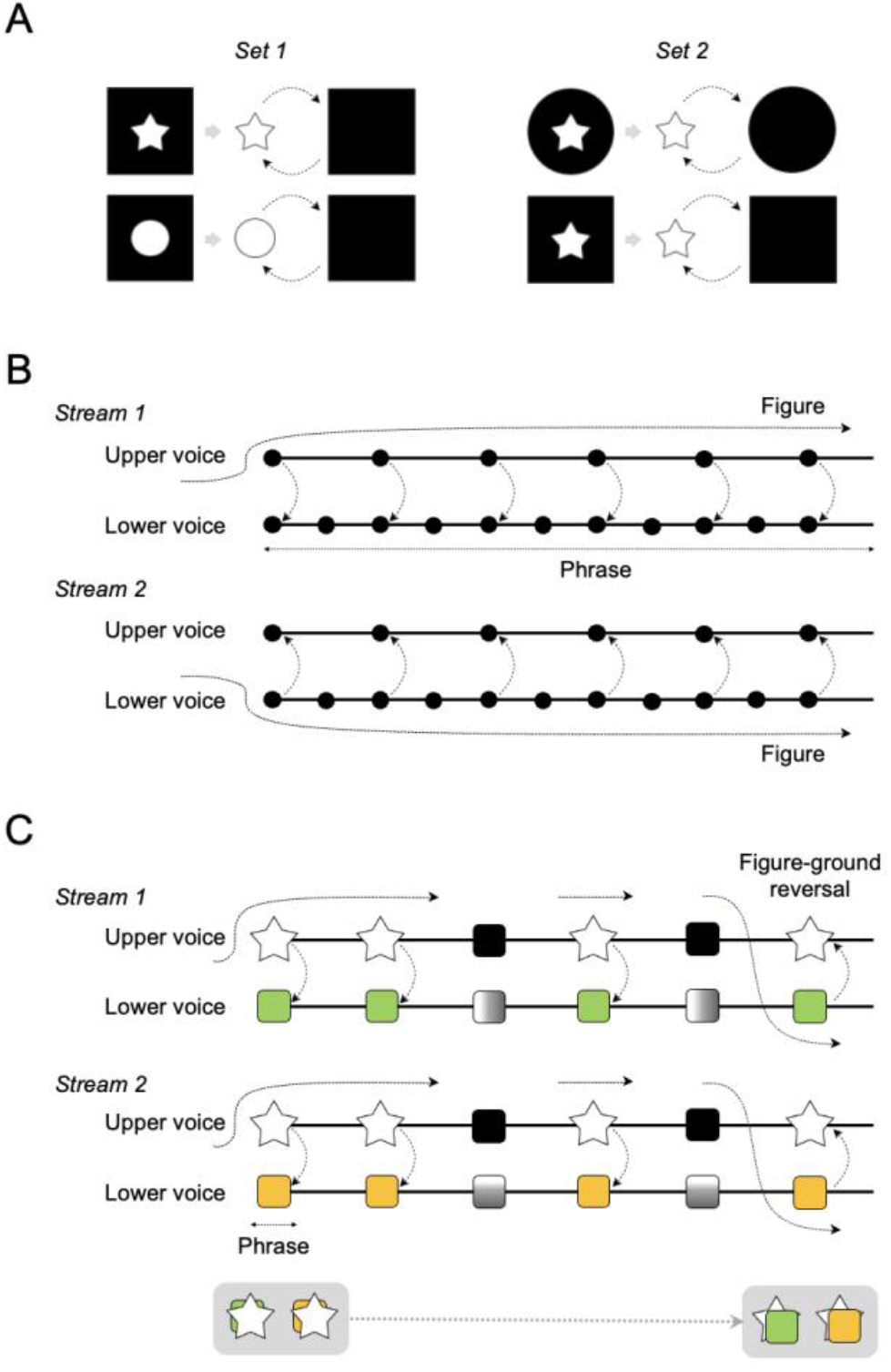
Figure–ground perception in auditory and visual stimuli. **(A)** Set 1 and 2 illustrate the examples of figure–ground perception in visual stimuli. In Set 1, both objects share the same black square background. However, the black squares can become figures, whereas the white star and circle are perceived as the ground. In Set 2, when the white stars form the figure perceptually, the black circle and square serve as the ground, and vice versa. Notably, the difference between the two objects lies in the shapes of the black circle and square. **(B)** In multivoiced music (Stream 1), the lower voice in a phrase serves as ground while attention is directed to the upper voice as figure. Conversely, in Stream 2, the roles are reversed. This model was adapted from Bigand et al., 2000^11^. **(C)** Drawing on Bigand’s model, we developed a figure–ground reversal model. The upper voice of a familiar melody (denoted by the white star) is heard as the figure, in particular through the repetition of phrases in a formal structure. The repetition of the same phrases (black square) affects the anticipation of the reappearance of the phrase. This anticipation allows a shift of attention toward the lower voice. Consequently, the relationship between figure and ground in the upper and lower voices may naturally switch. The appearacne of the white star in front of the orange/green box can be reversed.

**Fig. 2.**
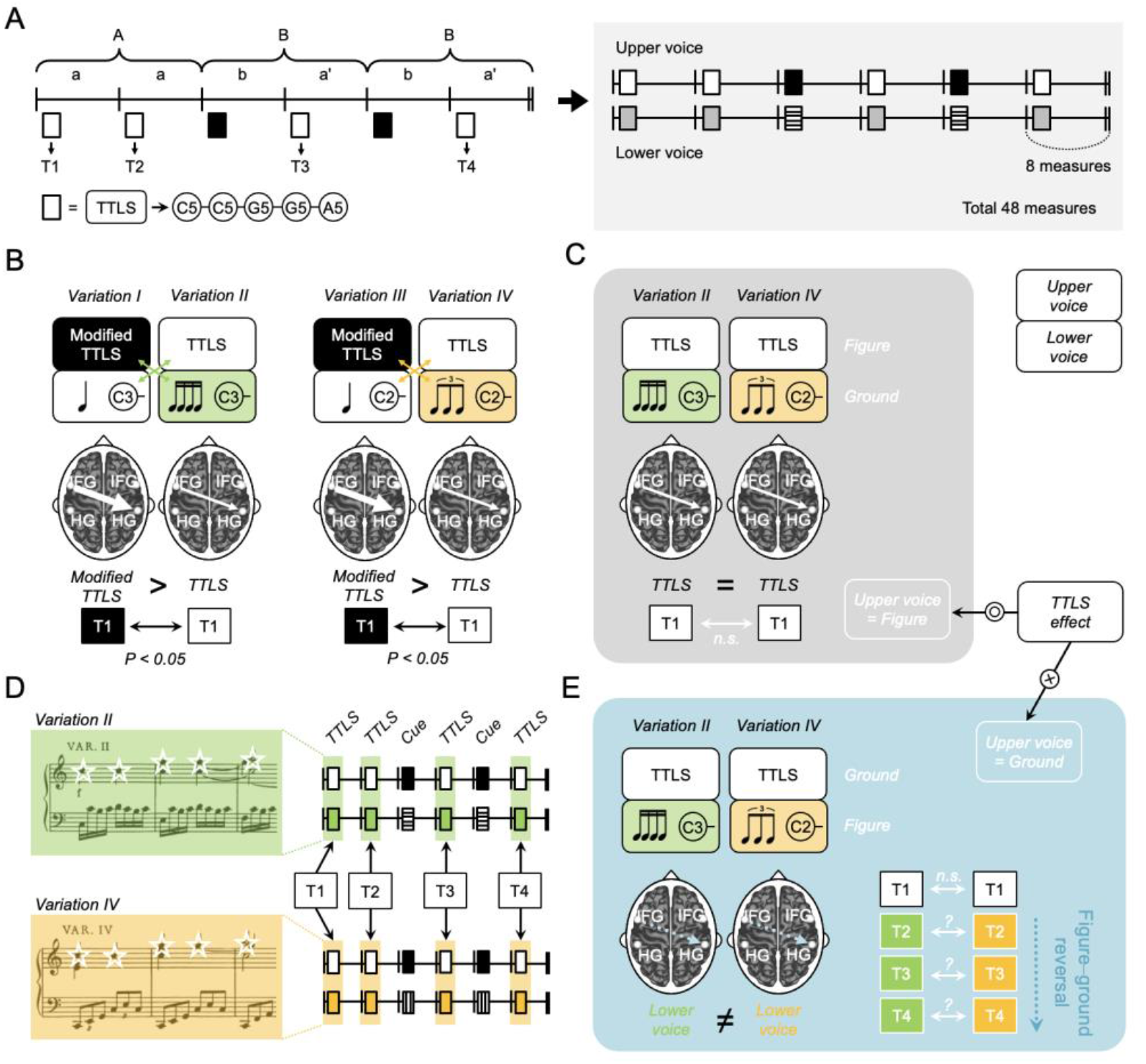
Figure–ground relationship between the upper and lower voices. **(A)** The left panel presents the entire structure of each of Mozart’s 12 Variations K. 265, including 48 measures of A (a + a) + B (b + a’) + B (b + a’) of a ternary form. The right panel presents two voices in the structure. The white squares denote the TTLS melody, C5-C5-G5-G5-A5, repeated four times per variation, while the black squares denote the cue phrase. **(B)** The melody of TTLS in the *Theme* is modified in *Variations I* and *III* but not in *Variations II* or *IV*. The rhythmic appearance is identical in each pair *Variations I* and *II* or *Variations III* and *IV*. The green cross shape and orange arrows denote the rhythm pattern shared between the voices of the two variations (Fig. S2). The frontotemporal connectivity from the left IFG to the right HG, depending on the presence of the TTLS melody, increased for the modified TTLS of novel information. The thickness of arrows between the left IFG and the right HG indicate the strength of the connectivity. **(C)** At T1, frontotemporal connectivity exhibits a consistent pattern between *Variations II* and *IV*, reflecting an effect of the TTLS melody. **(D)** The target phrases are highlighted as green and orange boxes, both containing the same white stars of the TTLS melody. The white and green/orange squares indicate repetitions of time windows among the 48 measures of each variation in the upper and lower voices, respectively. See Fig. S1 for details of the formation of the adapted scores, and the full scores. **(E)** When the figure–ground relationship is switched among T2–T4, unlike the case in T1, this means that the strength of the frontotemporal connectivity strength depends on the lower voices. The pattern of connectivity is unpredictable, because the two lower voices in *Variations II* and *IV* provide entirely novel information. See Fig. S2 for the definition of figure–ground reversal and the details of the hypothesis. TTLS, Twinkle Twinkle Little Star; T, target phrase

In our recent study using Mozart’s 12 Variations K. 265^10^, we observed changes in frontotemporal connectivity from the left Heschl’s gyrus (HG) to the left inferior frontal gyrus (IFG) in response to the presence or absence of the melody of the TTLS, C5-C5-G5-G5-A5. This response was consistent across target phrases lasting 2.1 s, equivalent to a five-quarter note length of C5-C5-G5-G5-A5 at the beginning of each variation in two pairs: *Variations I* and *II* and *Variations III* and *IV* (Fig. 2B). Thus, frontotemporal connectivity, specifically from the left IFG to the right HG, is not sensitive to lower voices having different rhythmic patterns but is responsive specifically to the TTLS melody (Fig. 2C). In our previous study^10^, we ruled out the effects of repetition by analyzing only the C5-C5-G5-G5-A5 at the beginning of each condition. If the relationship between the figure and ground reverses after repetition of the target phrases, the strength of the connectivity that is consistent between variations with the TTLS melody undergo changes. We developed a novel figure–ground reversal model (Fig. 1C) based on Bigand’s model^11^ on the relationship of figure and ground in a musical stream (Fig. 1B). The detection of the figure–ground reversal in the repeated target phrases fundamentally involves the processing of musical contexts built up by the target phrases that reiterate in each variation. In Mozart’s 12 Variations K. 265, we selected the two conditions of *Variations II* and *IV* (Fig. 2D) and reviewed how the frontotemporal connectivity that is consistent between two variations for the first target phrase in our previous study^10^ changes in target phrases (T1 to T4) containing the TTLS melody C5-C5-G5-G5-A5, repeated four times per variation. We hypothesized that, given the change between *Variations II* and *IV* in terms of frontotemporal connectivity observed in a target phrase, unlike in our previous results^4^, it could be caused by figure–ground reversal. Consequently, we developed two hypotheses: (1) If the frontotemporal connectivity for target phrases does not significantly differ between *Variations II* and *IV*, the same melody is processed as the figure, with the lower voice remaining the ground (Fig. 2C). (2) If there is a significant difference in the frontotemporal connectivity between the two variations, differences in the lower voices lead to a conversion of the lower voices to the figure, converting the upper voices to the ground (Fig. 2E). The property of the frontotemporal connectivity in the former, TTLS connectivity, would change in the latter, as the process focuses on the lower voices, which differ between variations, rather than on the upper voice of the same TTLS melody. Thus, in this study, we defined the neuroscientific phenomenon of figure–ground reversal as difference in frontotemporal connectivity that is observed between two variations sharing the TTLS melody but differing in the lower voices (Fig. S2). The difference in frontotemporal connectivity was tested for the same pairs of each target phrase between the two variations, respectively (Fig. S2C).

We estimated the effective connectivity from the left IFG to the right HG using linearized time-delayed mutual information (LTDMI) ^12^, a measure employed in our previous study^10^. LTDMI allows for estimating information transmission between two regional sources and observing the directionality in interhemispheric and interregional connectivity. We anticipated that the perceptual reversal between the figure of the melodic line and the ground of the lower voice line would elicit distinct responses within the repetitions of the target phrases in the musical structure (Fig. 2A). The dynamic interplay of brain networks between bilateral IFGs and HGs is instrumental in processing various musical components. Particularly, the left IFG plays a critical role in implicit learning of syntax in both language and music^13^, also contributing to memory contolling^14^ and attention^15^. Thus, the directionality of connection from the left IFG to the right HG efficiently explains the memory-based processing of input stimuli, including the aforementioned processes. This approach allowed us to explore how changes in effective connectivity within these regions correlate with perceptual shifts associated with repeated target phrases, shedding light on the intricate dynamics of melodic and lower voice processing.

While the central concept of this study revolves the shifting of attention between voices, the primary objective of this study was not to investigate connectivity changes through the deliberate manipulation of subjects’ attention to a particular voice or to assess the effects of attention. Instead, this study aimed to discern the musical scene in which the figure and the ground switch within the brain signals of individuals listening to the naturalistic stimuli of Mozart’s 12 Variations K. 265. During the MEG experiment, all subjects passively listened to the musical stimulus once, without any instructions to focus their attention on a specific voice or melody. We did not collect behavioral responses regarding respect to their attention shift toward the figure of the TTLS melody or the ground of the lower voice during the MEG recording or before and after the recording. If participants had been given any instructions concerning the figure–ground relationship of the voices before the presentation of the stimulus in the MEG recording for parallel behavioral recording, they would likely be inclined to focus on and select either the upper or lower voice while listening to the stimulus. This would invalidate our hypothesis concerning the natural switch in relationship between the figure and ground. Even where the behaviors are separately recorded, participants might not accurately remember the events of the individual target phrases in *Variations II* and *IV* following listening to the context from the theme to *Variation IV*. Moreover, as the participants were passively listening to music in the experiment, although the figure–reversal occurs, the participants may not recognize the moment. Thus, it would be impossible and fruitless to obtain valid results for our hypotheses using behavioral response. Our inquiries were based on the careful selection of repeated target phrases designed to elicit figure–ground reversal within a formal structure based on music theory. We constrained our hypotheses using the TTLS connectivity, which effectively controlled for other musical dynamics outside of the TTLS melody. Thus, our current hypotheses are supported by results from both music theoretical and neuroscientific analyses.

## Results

### LTDMI differences between two variations

Frontotemporal connectivity from the left IFG to the right HG was assessed using LTDMI^9^. We independently performed a Wilcoxon signed-rank test for each target period to confirm that the change in the figure–ground relationship between the two voices could be reflected in the frontotemporal connectivity from the left IFG to the right HG across variations. The difference in frontotemporal connectivity between *Variations II* and *IV* was calculated for four target phrases (T1–T4), each repeated four times per variation (Fig. 2). The difference between the two variations was significant only in T4 among all target phrases, as indicated by the Wilcoxon signed-rank test (Z = −2.112, *P* = 0.035; Fig. 3A). In T4, frontotemporal connectivity was enhanced in *Variation IV* compared with *Variation II*. However, significant differences were not observed in T1–T3 (*P* > 0.05 in all cases). In addition, we confirmed that, among 12 connections between the bilateral IFGs and HGs, the only significant result corresponded specially to frontotemporal connectivity from the left IFG to the right HG (Table S1). We observed a result that was close to the level of significance in temporofrontal connectivity from the right HG to the left IFG, in the opposite direction of frontotemporal connectivity (Right HG → Left IFG, Z = −1.843, *P* = 0.065; Fig. 3A and Table S1). This was not initially proposed by us in our hypothesis. The significance level (α) for the null hypothesis was independently tested for the individual target periods (T1–T4), since individual target phrases existed in completely different musical contexts before and after in the formal structure (Fig. 2A).

**Fig. 3.**
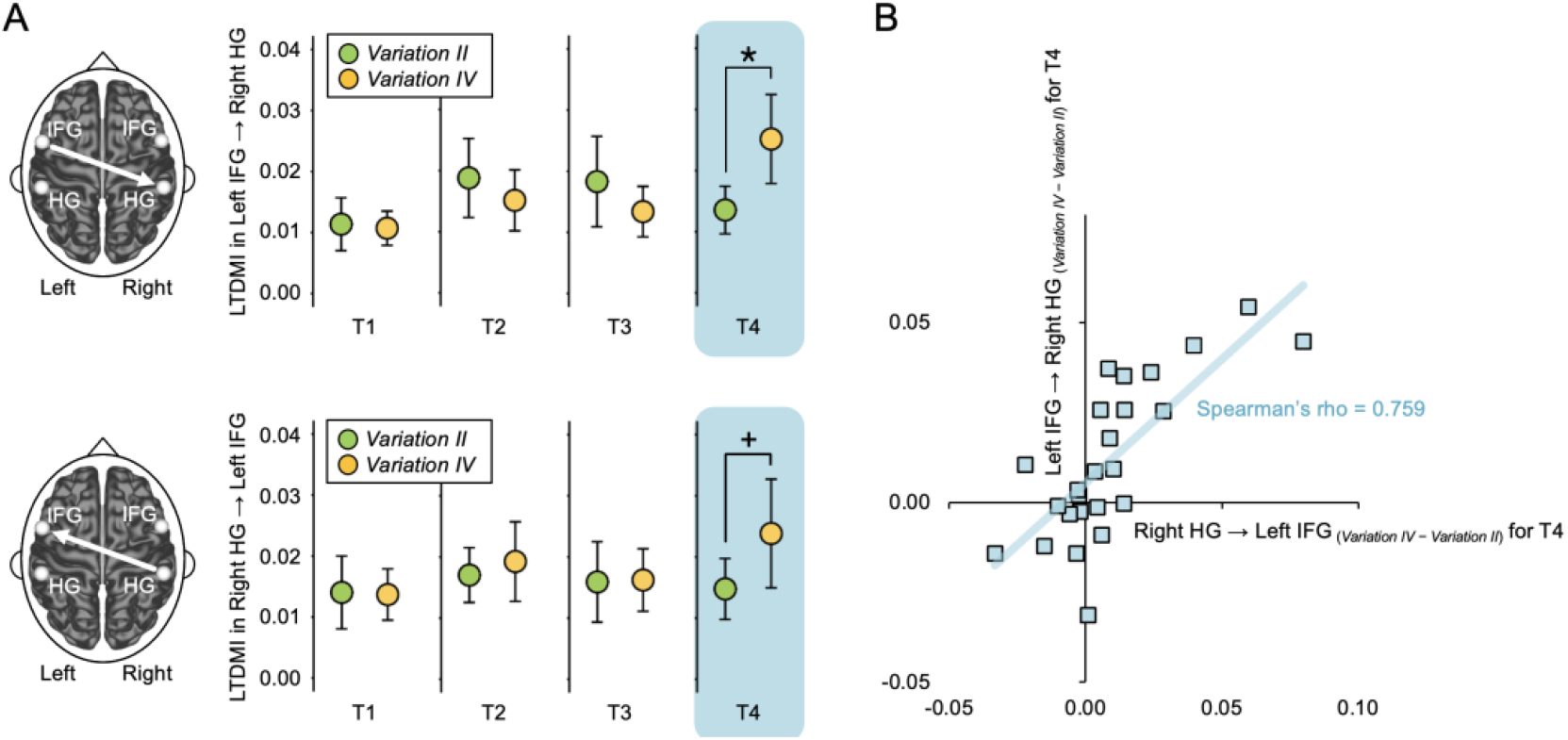
Change in connectivity through figure–ground reversal. **(A)** Frontotemporal connectivity of *Variation IV* was significantly enhanced compared with that of *Variation II* only during T4 (Wilcoxon signed-rank test, Z = −2.112, *P* = 0.035,). The temporofrontal connectivity (right HG ⟶ left IFG) of *Variation IV* was also enhanced relative to that of *Variation II* only at T4, although this did not reach the level of statistical significance (Wilcoxon signed-rank test, Z = −1.843, *P* = 0.065). For both frontotemporal and temporofrontal connectivity, there were no significant differences from T1 to T3 (*P* > 0.05 in all cases; Wilcoxon signed-rank test). Error bar denotes the standard error mean. *, *P* < 0.05. **(B)** Frontotemporal connectivity (Left IFG → Right HG) was strongly positively correlated with the temporofrontal connectivity (Right HG → Left IFG) only during T4 (Table S2). There was a significant correlation between Left IFG → Right HG _(*Variation IV*– *Variation II*)_ and Right HG → Left IFG _(*Variation IV*– *Variation II*)_ at T4 (Spearman correlation, Spearman’s rho = 0.759, Bonferroni-corrected *P* = 0.0001). “*Variation IV*– *Variation II*” denotes a difference between *Variation IV* and *Variation II* for the LTDMI value. LTDMI, linearized time delayed mutual information; HG, Heschl’s gyrus; IFG, inferior frontal gyrus; T, target phrase

We defined the figure–ground reversal phenomenon as the difference in frontotemporal connectivity between two variations (Fig. S2). As expected, a difference between the variations was observed in T4. The LTDMI value was higher in *Variation IV* than in *Variation II*. If this is caused by the difference in musical features in the lower voices of both variations, we were not able to determine what specific (novel) information triggered the observed connectivity attenuation and enhancement in *Variation II* and in *Variation IV*, respectively. The musical features in the lower voice for *Variation II* would affect the enhancement of the frontotemporal connectivity less. However, if connectivity attenuation in *Variation II* indicates a consistency of the LTDMI value due to the effect of TTLS melody, the figure–ground reversal would not occur in *Variation II*. Thus, we interpret the connectivity reduction in *Variation II* for T4 as caused by the properties of the lower voices in *Variation II*, which differed from those in *Variation IV* (Fig. 4A) or by the effect of the TTLS melody, which was not switched in *Variation II* (Fig. 4B).

**Fig. 4.**
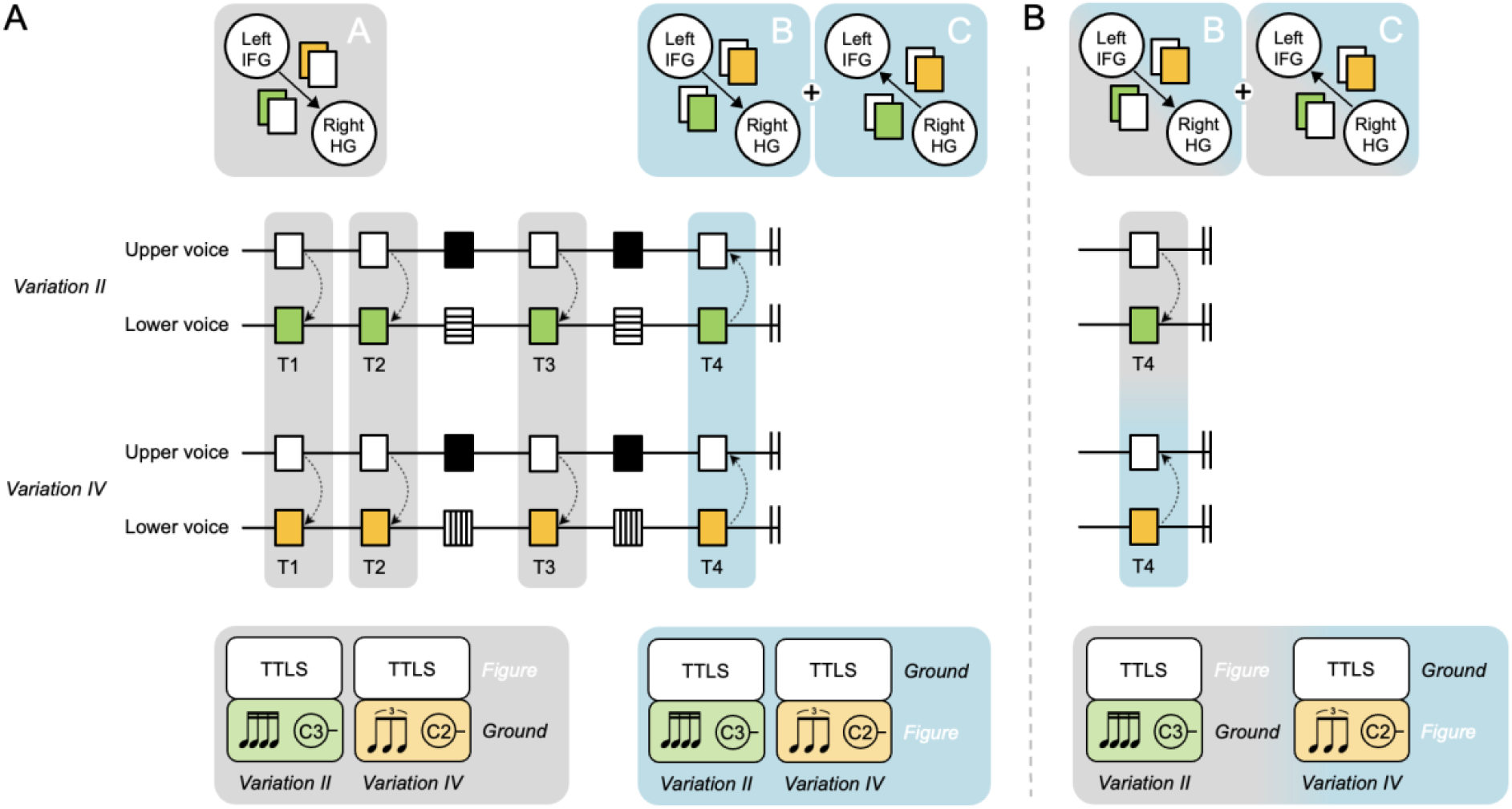
Directional connectivity in relation to the figure–ground relationship. **(A)** The diagram shows how the connectivity between the left IFG and the right HG alters from T1 to T4 with the repetitions of the same passages for each variation. Repeated and cue phrases can affect the anticipation of the TTLS melody (white square). Frontotemporal connectivity from the left IFG to the right HG, the TTLS connectivity, is consistent from T1 to T3, focusing on the figure of the upper voice. “A” could indicate, “I know this song.” However, at T4, A different frontotemporal connectivity pattern is exhibited between *Variations II* and *IV*. This indicates that frontotemporal connectivity is not related to the TTLS melody, but the lower voice switched as in the figure, reflecting that frontotemporal connectivity is no longer the TTLS connectivity. “B” could indicate, “I know that there are different elements apart from the song.” For T4, the temporofrontal connectivity, the opposite direction of the frontotemporal connectivity, also exhibits a consistent pattern. “C” might indicate, “I know how the elements differ.” **(B)** However, if the attenuation of connectivity in *Variation II* indicates the consistency of the value of the LTDMI value due to the effect of the TTLS melody, the figure–ground reversal would not occur in *Variation II*. The relatively low LTDMI value in T4 of *Variation II* may indicate that the figure–ground reversal is less seen in *Variation II*. White squares denote phrases of the upper voice of the TTLS melody common to the two variations. Green and orange squares represent lower voices that differ between the variations. Black squares indicate cue phrases. HG, Heschl’s gyrus; IFG, inferior frontal gyrus; T, target phrase; TTLS, Twinkle Twinkle Little Star

### Correlation between frontotemporal and temporofrontal connectivity

Correlation analyses were conducted to confirm (1) whether a similar pattern between frontotemporal and temporofrontal connectivity refers to bidirectional information transmission between the left IFG and the right HG and (2) whether a similar pattern is only specialized in the temporofrontal connectivity (Right HG → Left IFG) among 12 connections between the bilateral IFGs and HGs, which are key areas for the music process. To perform this estimation, we first computed the difference values between *Variations IV* and *Variation II* for the LTDMI values in 12 connections between the bilateral IFGs and HGs for all target phrases of T1 to T4. Next, we estimated the correlation between the frontotemporal connectivity difference value (Left IFG → Right HG _(*Variation IV*– *Variation II*)_), with 11 other connectivity difference values. In the Spearman correlation test result, a significant correlation was only observed between Left IFG → Right HG _(*Variation IV*– *Variation II*)_ and Right HG → Left IFG _(*Variation IV*– *Variation II*)_ for T4, among 44 combinations of 11 connections × 4 target phrases (Spearman’s rho = 0.759, Bonferroni-corrected *P* = 0.0001; Fig. 3B and Table S2). The frontotemporal connectivity (Left IFG → Right HG) was strongly positively correlated with the temporofrontal connectivity (Right HG → Left IFG).

## Discussion

Difference in frontotemporal connectivity from the left IFG to the right HG between the two variations was only observed in the final target phrase of T4, without differences in the preceding three phrases. According to our hypothesis, an examination of the disparity in the LTDMI between the two variations suggests that the TTLS melody of the upper voice underwent a reversal, becoming the ground in T4, but remained the figure in T1–T3 (Fig. 3A). This indicates that subjects may have been particularly attentive to the TTLS melody up to T3, but in T4, a natural shift occurred, leading them to focus more on the lower voice due to habituation from repeated listening. Each variation included cue phrases like “Up above the world…,” predicting the recurrence of the TTLS melody (Fig. 2A and S1). T4 was introduced after the second iteration of the cue phrase (Fig. 2A), helping subjects anticipate the reappearance of the TTLS melody. The predictability of the upper voice through repetition might have facilitated participants’ recognition of the lower voice^7^. In addition, we found a consistent pattern in that the mean LTDMI value was enhanced in *Variation IV* relative to *Variation II* in the temporofrontal connectivity, which was the opposite direction to the frontotemporal connectivity (Fig. 3A). In our previous study^10^, unidirectional information transmission from the left IFG to the right HG, showing frontotemporal connectivity, was associated with the detection of the TTLS melody. Thus, we were not able to conclude whether the temporofrontal connectivity pattern, consistent from T1 to T3, referred to the same process in the frontotemporal connectivity, of which the TTLS melody is the figure (Fig. 4). The change in temporofrontal connectivity was observed in T4. Furthermore, in T4, a similar pattern with frontotemporal connectivity is connected to positive correlation (Fig. 3B and Table S2). Finally, our data indicate that the bidirectional connectivity between the left IFG and the right HG in T4 collaborates in the processing of figure–ground reversal between the TTLS melody and the lower voice. Frontotemporal connectivity from T1 to T3 could indicate, “I know this song.” In T4, the frontotemporal connectivity might indicate, “I know that there are different elements apart from the song,” and the temporofrontal connectivity might indicate, “I know how the elements differ” (Fig. 4).

Difference in frontotemporal connectivity from the left IFG to the right HG between the two variations, was observed only in the final target phrase of T4, without differences in the preceding three phrases. According to our hypothesis, an examination of the disparity in the LTDMI between the two variations suggests that the TTLS melody of the upper voice underwent a reversal, becoming the ground in T4, but remained the figure in T1–T3 (Fig. 3A). This indicates that subjects may have been particularly attentive to the TTLS melody up to T3, but in T4, a natural shift occurred, leading them to focus more on the lower voice due to habituation from repeated listening. Each variation included cue phrases like “Up above the world…,” predicting the recurrence of the TTLS melody (Fig. 2A and S1). T4 was introduced after the second iteration of the cue phrase (Fig. 2A), helping subjects anticipate the reappearance of the TTLS melody. The predictability of the upper voice through repetition might have facilitated participants’ recognition of the lower voice^7^. Additionally, we found the consistent pattern that the mean LTDMI value was enhanced in *Variation IV* compared to *Variation II* in temporofrontal connectivity, which is the opposite direction of the frontotemporal connectivity (Fig. 3A). In our previous study^10^, the unidirectional information transmission from the left IFG to the right HG, referring to frontotemporal connectivity, was associated with the detection of the TTLS melody. Thus, we could not conclude whether the temporofrontal connectivity pattern, consistent from T1 to T3, refers to the same process of in the frontotemporal connectivity, which the TTLS melody is the figure (Fig. 4). The change in temporofrontal connectivity was observed in T4. Furthermore, in T4, the similar patter with frontotemporal connectivity is connected to a positive correlation (Fig. 3B and Table S2). Finally, our data show that the bidirectional connectivity between the left IFG and the right HG in T4 collaborates for the processing of figure–ground reversal between the TTLS melody and the lower voice. The frontotemporal connectivity from T1 to T3 might indicate, “I know this song”. In T4, the frontotemporal connectivity might indicate, “I know that there are different elements apart from the song”, and the temporofrontal connectivity might indicate, “I know how the elements differ” (Fig. 4).

Top-down processing of a familiar melody can significantly influence the figure–ground relationship^16,17^. The left IFG is crucial in processing familiarity^18^ and plays a key role in music-syntactic processes, indicating implicit learning^13^ and contributing to memory retrieval^14,19^. In addition, the left IFG is involved in differentiating between melody and accompaniment^20^. On the other hand, the right HG predominantly processes tone deviance^21,22^ and the segregation of auditory streams^23^. The right auditory cortex has been the dominant site for music processing^24^. Accordingly, the directional information flows in frontotemporal connectivity could explain how the left IFG and the right HG collaborate in processing target phrases. The IFG and HG are pivotal areas for music perception, and their connectivity is discussed in relation to syntax processes^25-27^, categorization^28^, and the working memory of melody process^29^. Taking into account the roles of frontotemporal connectivity that were linked to the TTLS melody-based process and temporofrontal connectivity linked to the sensory-based process, this heightened connectivity could indicate a dual process: extracting the novel lower voice in the target phrase including the familiar TTLS melody, and dissecting the components within the novel lower voice. The temporofrontal connectivity supports the frontotemporal connectivity’s process. Thus, the figure–ground reversal recruited the bidirectional connectivity of Left IFG → Right HG and its opposite direction, Right HG → Left IFG, referring to both a top-down process based on knowledge of the TTLS melody and a bottom-up process based on new information on voices accumulated while sequentially listening to target and cue phrases in each variation^30,31^.

Familiarity is critical for explaining the figure–ground experiment^16,32,33^. The target phrases in our stimuli also involved the TTLS song, used in studies^34-37^ related to perception of familiar melodies. Familiarity would be naturally implied in the theme and all variations, considering that Mozart’s 12 Variations K. 265 is based on a theme of the TTLS melody. Logically, the familiarity implied in the TTLS melody is influential on figure–ground perception and its reversal. However, in our previous study^10^ focusing on the TTLS melody, we could not directly prove the effect of familiarity on frontotemporal connectivity. Frontotemporal connectivity was the connectivity that changed with the presence and absence of the TTLS melody in our previous study^9^. In our present study, the TTLS melody appearing repeatedly was the same in *Variation II* and *Variation IV*. The effects of familiarity are consistent with both *Variations II* and *IV*. TTLS connectivity was applied to prove the figure– ground reversal in our present study, throwing the following results: (1) Consistent TTLS connectivity between *Variations II* and *IV* from T1 to T3 indicates that the TTLS melody in the target phrases served as the figure (Fig. 2C). (2) Inconsistent TTLS connectivity between *Variations II* and *IV* in T4 indicates that the TTLS melody in the target phrases functioned as the ground, while the lower voice became the figure (Fig. 2E).

Research on figure–ground reversal originated in visual studies, notably using the classic Rubin vase-faces model^17,38^. Monkey data have highlighted the involvement of the primary visual cortex in neuronal activities in response to figure–ground perception of visual stimuli^39-42^. Visual figure–ground discrimination as well as behavior, has been explored in various animals such as mice^43^, flies^44^, and honey bees^45^. Notably, figure–ground processing in visual objects differs between anesthetized and awake animals^46^. In the present study, behavioral responses to differences between T1, T2, T3, and T4 were not measured, because task performance while listening to music did not align with the objective of this study. Brain responses were solely measured in the context of figure–ground reversal using real music, without controlling for attention. While previous studies have selectively manipulated stimuli or controlled subjects’ attention to focus on specific auditory streams^20,47-52^, attention’s critical role in figure–ground perception has been acknowledged^53^. Despite our results, it remains unclear whether both figure and ground were processed attentively or pre-attentively, given the absence of intentional attention control in our study. Artificially manipulated stimuli, while useful in some cases^54,55^, deviate from the complex and dynamic nature of real musical experiences. Music listeners can automatically process information such as syntactic errors and tone deviations even without intentional attention^56,57^. Although listeners may focus on a particular voice while listening to music, the ever-changing flow of music can result in such brief perceptible moments that the figure–ground relationship continuously shifts without listeners consciously realizing. Our results successfully captured the moment when the figure–ground relationship between the upper and lower voices switched, as evidenced by the difference in frontotemporal connectivity for repeated phrases in the two variations, as well as by the correlation between frontotemporal and temporofrontal connectivity.

For auditory stimuli, figure–ground perception is not exclusive to humans; it has also been observed in animals, in studies on figure–ground processes investigated through noise and sound. Extracellular recordings demonstrated increased multiunit activities in the auditory area^58^. Further, auditory areas have been the focus of previous human studies on stream segregation through selective attention and figure–ground segregation^47,48,50,52^. In fact, numerous studies on auditory figure–ground segregation were based on the concept of auditory scene analysis^59^ and grouping^1^, using tasks that discriminate a sound pattern as a figure from a ground of tone and chord sequences with irregularities in spectral and temporal properties^54,55,60^. Regions involved in the discrimination of figure–ground segregation reported in fMRI and EEG studies^54,55,60^ comprise the primary auditory area, superior temporal sulcus, superior temporal gyrus, intraparietal sulcus, medial/superior frontal gyrus, and cingulate cortex. In addition to these findings, the processing of multiple voices in this study involved the IFG and the HG, both of which exhibited information transmission. This difference in involvement may be attributed to the nature of our stimuli, which comprised real music containing contextual richness rather than edited notes or sequences. Furthermore, the participants in this study passively listened to Mozart’s 12 Variations K. 265 without being asked to concentrate selectively on specific musical parts. The involvement of the left IFG and right HG might reflect the entire process of naturally grasping, comparing, and understanding the voices of target phrases in the context, rather than simply listening to a sound. Therefore, bilateral information transmission that involves both frontotemporal and temporofrontal connectivity encompass all activities of discriminating and dissecting target phrases with the TTLS melody repeated within the real music of Mozart’s 12 Variations K. 265. This includes segregating the lower voice from a stream containing the TTLS melody and perceiving the figure–ground reversal of the TTLS melody and its lower voices in the musical context of the repeated target and cue phrases in Mozart’s 12 Variations K. 265.

Naturalistic stimuli have been used to examine various topics^61-63^. In studies of the concepts of emotion^64,65^, melodic expectation^66^, temporal aspects of rhythm and beat^67,68^, and familiarity^69^, multiple naturalistic pieces have been used as musical stimuli. However, in this present study, it would not be appropriate to conduct examination across multiple stimuli for following reasons. Although we could select as musical stimuli multiple pieces with having a familiar melody, it is essential to clarify whether they would have a level of familiarity similar to the TTLS melody. In addition, the musical context, including target and cue phrases and their repetitions, as well as the consistency of tonality, harmonic structure, timbre, tempo, and texture, should all be taken into account. Moreover, due to the inherent characteristics of naturalistic stimuli that cannot be perfectly controlled, there may be complex variables that are capable of eliciting subtle differences in the results. Thus, study using naturalistic stimulus could examine their hypotheses, including motif, musical features, timbre, and depression, based on a single piece^70-73^.

The figure–ground examination in this study was performed based on the TTLS connectivity that was identified in our previous study^10^. A fair test of a hypothesis should include, at a minimum, evaluations having a range of different stimuli. However, our hypothesis was created for the melody of TTLS and its perceptual reversal using Mozart’s 12 Variations K. 265. The figure–ground reversal phenomenon can be commonly observed in various fields of audiation and vision, as noted. The phenomenon of a switched relationship between the elements of an object can be simply observed and explained though the comparison of related objects. However, naturalistic music has its own narrative, which can be described by its context. Indeed, although our hypothesis focused on *Variations II* and *IV*, the processes included the context from the theme to *Variation IV*, including *Variations I* and *III*, where the melody is a modification of the TTLS melody (see Fig. 2). Individual variations were structured independently in a ternary form, with a pause (rest) between them. All of the participants listened to the overall context from the theme to *Variation IV* sequentially. Furthermore, our hypothesis did not concern a perceptually dominant melody in the highest voice but the perceptual reversal in multiple voices, including the lower voices, showing different rhythmic patterns. Thus, the connectivity reflects complicated processes not only for the target phrases of 2.1 s without the context but also for each 2.1 s target phrases in the context of the theme and variations, leading up to the target phrase. This is why we examined figure–ground reversal using naturalistic music. This approach, however, may form a critical shortcoming of our study, compared to measurements using artificially composed stimuli. Nevertheless, our results completely reflected our ubiquitous experiences of individual subjects.

In addition to its use of a single piece of naturalistic music, our study had other limitations. While nonmusicians can perceive the figure separately from the ground^54^, they tend to focus more on the upper melody than on a lower voice^6^. In contrast, musicians exhibit greater sensitivity to voice perception and superior ability in distinguishing voices^4,51^. Music training significantly influences selective attention^49^. We did not recruit musicians as participants to explain the figure–ground reversal in terms of basic musical ability. We consider that recruiting nonmusicians could have impacted our results as follows: (1) In terms of frontotemporal connectivity, if the LTDMI during T4 in *Variation II* reflects both the processing of lower voice features and the absence of figure–ground reversal, the outcome could differ for musicians who have greater sensitivity. (2) If musicians were participants, temporofrontal connectivity could show a statistically significant distinction (*P* < 0.05), since recognition of different elements in the lower voices (via temporofrontal connectivity) would involve a more complex process of cognition. Some nonmusicians might process this successfully, while others might not. Therefore, future studies should verify these results with musicians. In addition, this study concentrated on two regions, the left IFG and the right HG. Our findings should be verified at the whole-brain level. Nonetheless, using Mozart’s 12 Variations K. 265 was a compelling and valuable endeavor in unraveling the fundamental neural processes associated with processing real music featuring multiple voices and understanding the human experience of music. Our data effectively illustrated how the brain can dissect voices from the multidimensional structures of continuously changing music and reconstruct them through a momentary switch in figure–ground processes.

## Materials and Methods

### Participants

Participants comprised 25 healthy individuals, all nonmusicians, 15 women and 10 men with a mean age of 26.8 ± 3.4 years old. None had received formal musical training. All participants were right-handed, with a mean Edinburg Handedness coefficient of 95.7 ± 7.1. The study adhered to the principles of the Declaration of Helsinki and received approval from the Institutional Review Board of the Clinical Research Institute at Seoul National University Hospital (IRB No. C-1003-015-311). The research procedures were conducted in accordance with relevant guidelines and regulations, and all subjects provided informed consent, adhering to ethical guidelines.

### Stimuli

Mozart’s K. 265 consists of the theme and twelve variations on “Ah! vous dirai-je Maman”. This study focused on *Variation II* and *Variation IV*, with the same TTLS melody. *Variations II* and *IV* share the same TTLS melody but differs in the lower parts, transforming through rhythmic changes to 8th note triplets and 16th notes (semiquavers), respectively. The tonality and harmonic structure are the same for both variations (Fig. 2A). Each variation was based on the ternary form of A (a + a) + B (b + a’) + B (b + a’). Phrases including the “C5-C5-G5-G5-A5” melody were repeated four times in each variation (Fig. 2A).

### MEG recording

In a magnetically shielded room, the participants listened to Mozart’s K. 265 while watching a silent movie clip (Love Actually, 2003, Universal Pictures, USA) for approximately 5 min. Musical stimuli were generated using STIM^2TM^ (Neuroscan, Charlotte, NC, USA) and presented binaurally at 100 dB through MEG-compatible earphones (Tip-300, Nicolet, Madison, WI, USA). MEG signals were recorded using a 306-channel whole-head MEG system (Elekta Neuromag Vector ViewTM, Helsinki, Finland) with a sampling frequency of 1,000 Hz and a bandpass filter of 0.1–200 Hz.

### MEG analysis

#### Source analysis

The MEG source signals of epochs from −100 ms to 2,100 ms after the onset of each condition for four regional sources of bilateral HGs and IFGs, bandpass filtered at 14–30 Hz, were extracted using BESA 5.1.8.10 (MEGIS Software GmbH, Gräfelfing, Germany) after electrooculograms, electrocardiograms, and muscle artifacts were removed. Standard Talairach coordinates (x, y, and z in millimeters) for bilateral HGs (transverse, BA 41, BA 42) and IFGs (triangular part, BA 45) across participants were adapted from previous research^10^. These coordinates were as follows: left HG (−53.5, −30.5, and 12.6), right HG (55.4, −30.5, and 12.6), left IFG (−55.5, 11.7, and 20.6), and right IFG (53.5, 12.7, and 20.6).

#### Time window

*Variations II* and *IV* were chosen as the two conditions for estimating connectivity differences, including the identical TTLS melody. Time windows were 2,100 ms long, incorporating the “C5-C5-G5-G5-A5” melody, as based on a previous study^10^. Time windows of 2,100 ms appeared four times per variation (Fig. 2D and Fig. S1). The lower voices accompanied by the “C5-C5-G5-G5-A5” melody differed between the two variations (Fig. 2). There was only one trial for each target phrase because the stimulus was real music. Each individual subject’s data included the data of four target phrases for the two variations.

#### LTDMI analysis

While our primary focus was the regional connection from the left IFG to the right HG, we verified our results for all connections among the bilateral IFGs and HGs. The effective connectivity for the 12 connections between regional sources of the bilateral HGs and IFGs was estimated using MATLAB 7.7.0.471 (Math Works Inc., Natick, MA, USA) (see also Table S1 for individual LTDMI values calculated for 12 connections). LTDMI was used to estimate information transmission between two time series based on mutual information (MI) ^12,25^. For each subject, the mean LTDMI for the 2,100-ms epoch was calculated for each of 4 target phrases (T1, T2, T3, and T4) × 2 variations (*Variations II* and *IV)*. Statistical comparisons of mean LTDMI values for *Variations II* and *IV* were performed using SPSS 21.0 software (IBM, Armonk, NY, USA). Considering different contexts in each target phrase, we conducted the non-parametric *Wilcoxon signed-rank test* due to the non-Gaussian distribution of LTDMI data. In each case, the significance level (α) for rejecting the null hypothesis (H0, indicating no difference between *Variation II* and *Variation IV* in the mean LTDMI values), was 0.05. In addition, in the nonparametric Spearman correlation test for each pair between the frontotemporal connectivity difference value (Left IFG → Right HG _(*Variation IV*– *Variation II*)_) and the other 11 connectivity difference values, except of Left IFG → Right HG among twelve connections between the bilateral IFGs and HGs, the Type I errors that were caused by multiple comparisons among the 11 connection pairs in the Spearman correlation test were adjusted by the Bonferroni test.

## Availability of Data and Materials

The datasets used and/or analyzed during the current study are available from the corresponding author on reasonable request.

## Acknowledgment

We sincerely appreciate Ji Hyang Nam for technically supporting MEG data acquisition.

## Author contributions

C.H.K.: conceptualization, funding, formal analysis, visualization, writing of original draft, and review and editing of the manuscript; J. S.: advice about music theory, and review and editing of the manuscript; C.K.C.; supervision, funding, and review and editing of the manuscript.

