## Supplementary figures and tables for "Bidirectional connectivity changes in the frontotemporal areas reflect figure–ground reversal in multivoiced music"

### Supplementary information



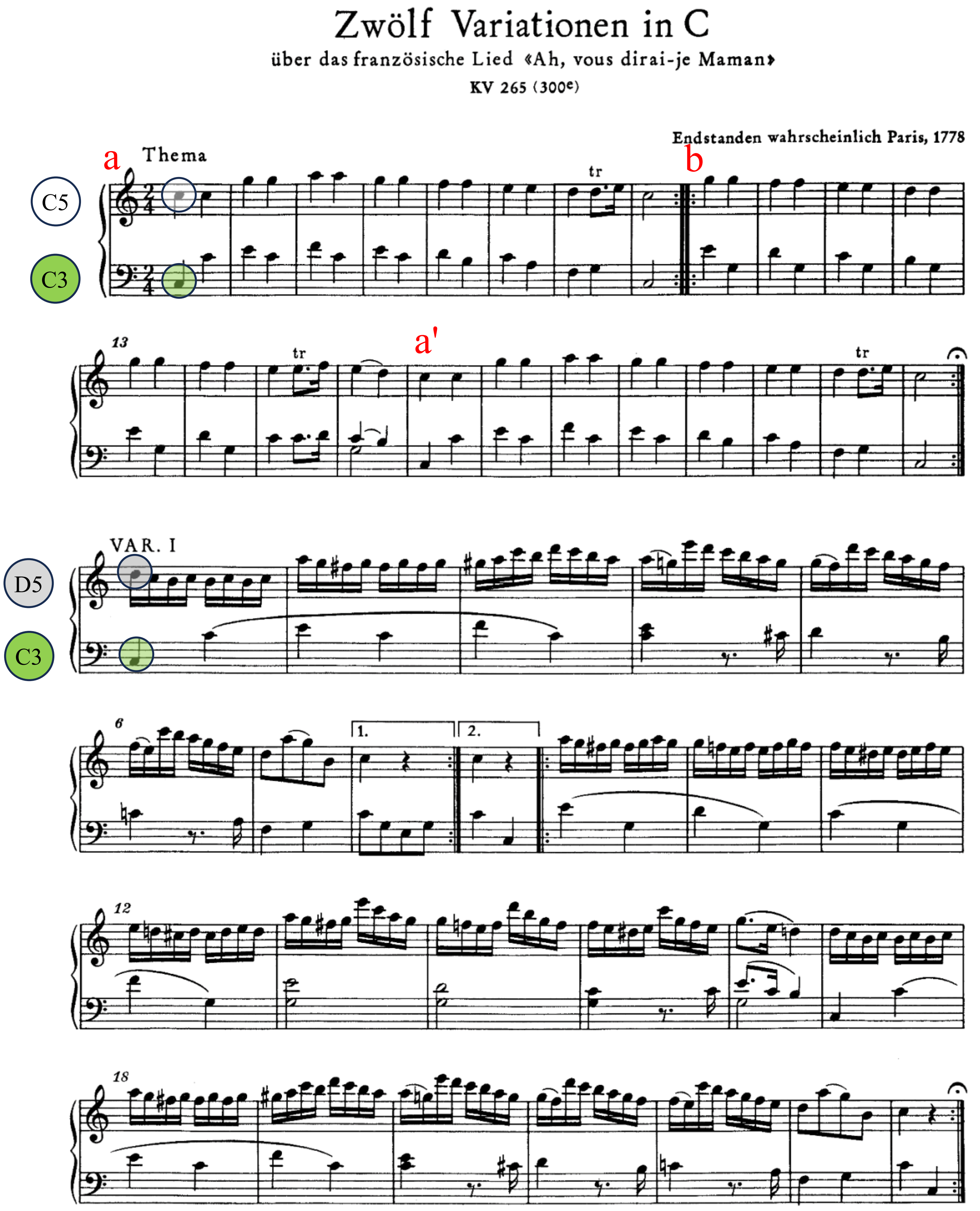

#### VAR. II

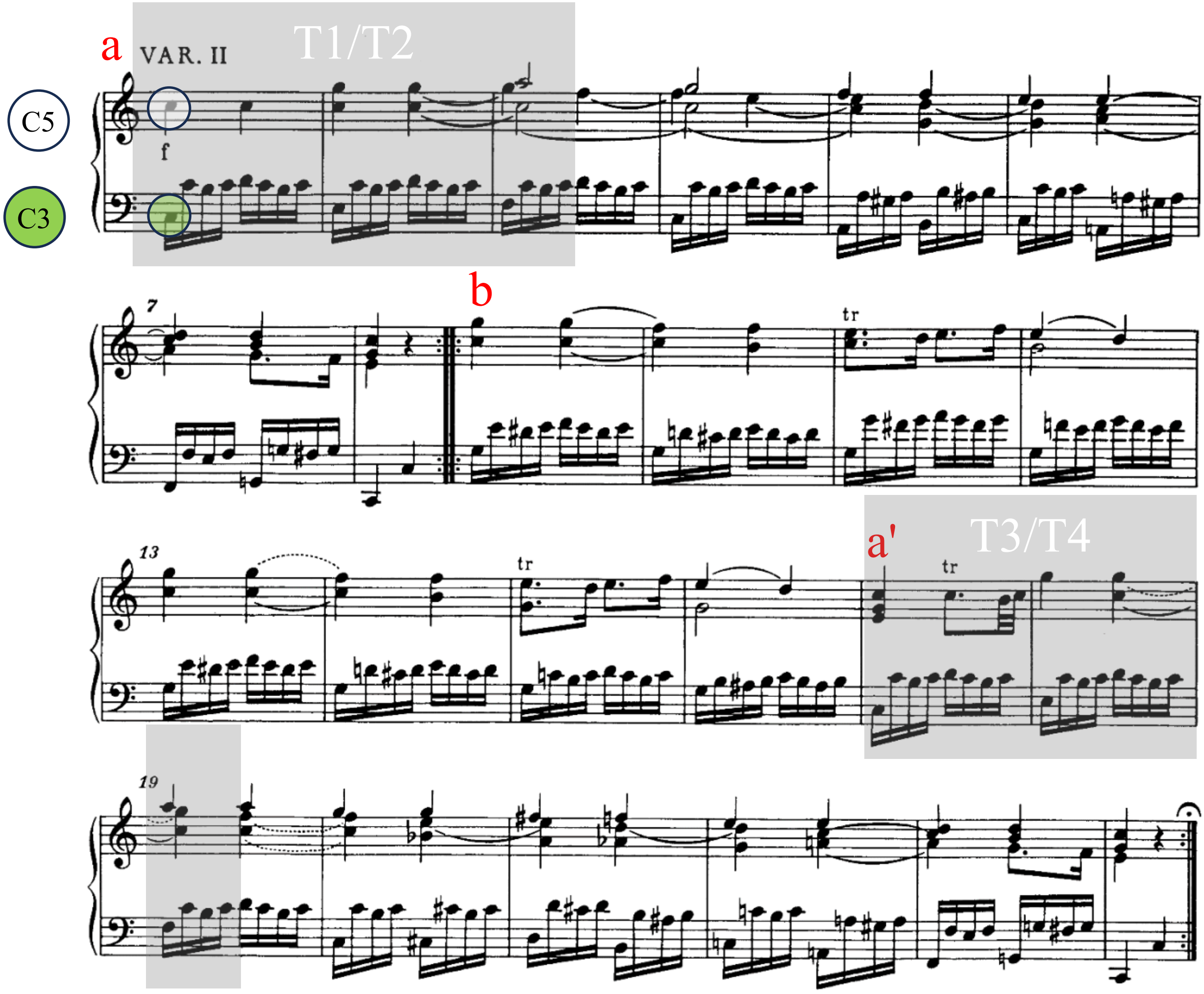


#### VAR. III

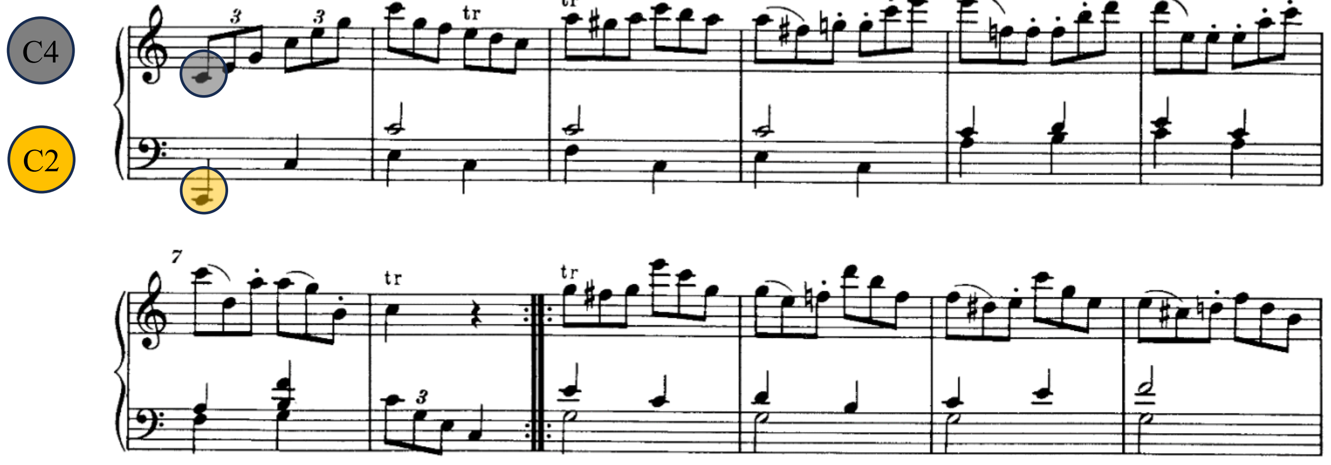

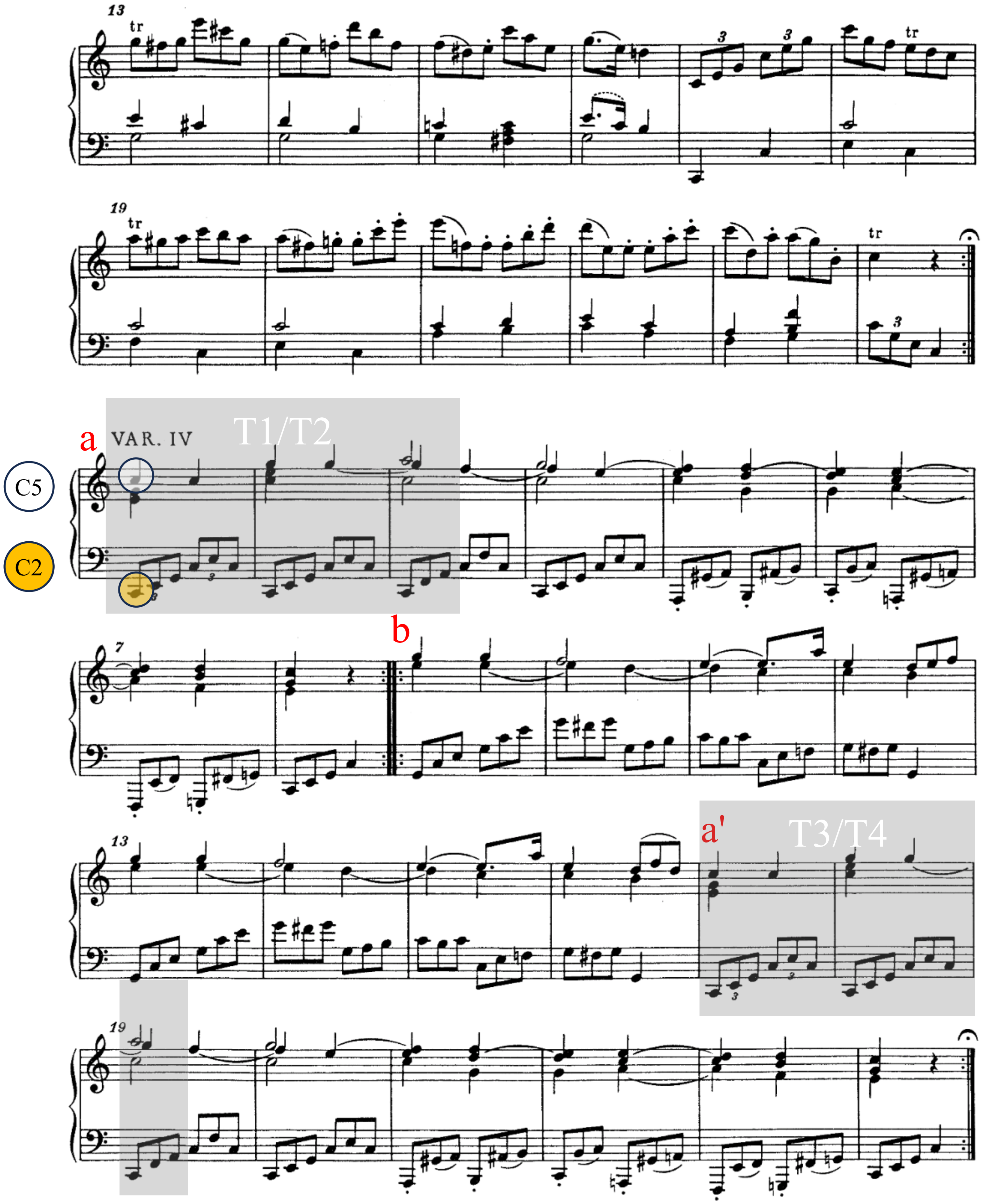
VAR. IV
T1/T2
a
b
a'
T3/T4
C5
C2

**Fig. S1. Musical stimulus.** Musical score of the theme to *Variation IV* in Mozart's 12 Variations on "Ah, vous dirai-je maman" K. 265 (Barenreiter edition) was adapted from *NMA Online: Neue Mozart-Ausgabe: Digitized Version* (https://dme.mozarteum.at/DME/nma/nmapub_srch.php?l=2). Each movement is based on the ternary form of A (a + a) + B (b + a') + B (b + a'). The time windows of T1, T2, T3, and T4 in *Variations II* and *IV* are marked by gray shaded boxes. Section "b" of each variation on the score includes cue phrases marked as black squares in Fig. 2. Pitch notations on white, green, orange, and gray circles at the beginning of each variation indicate the first notes of the upper and lower voices in each time window.

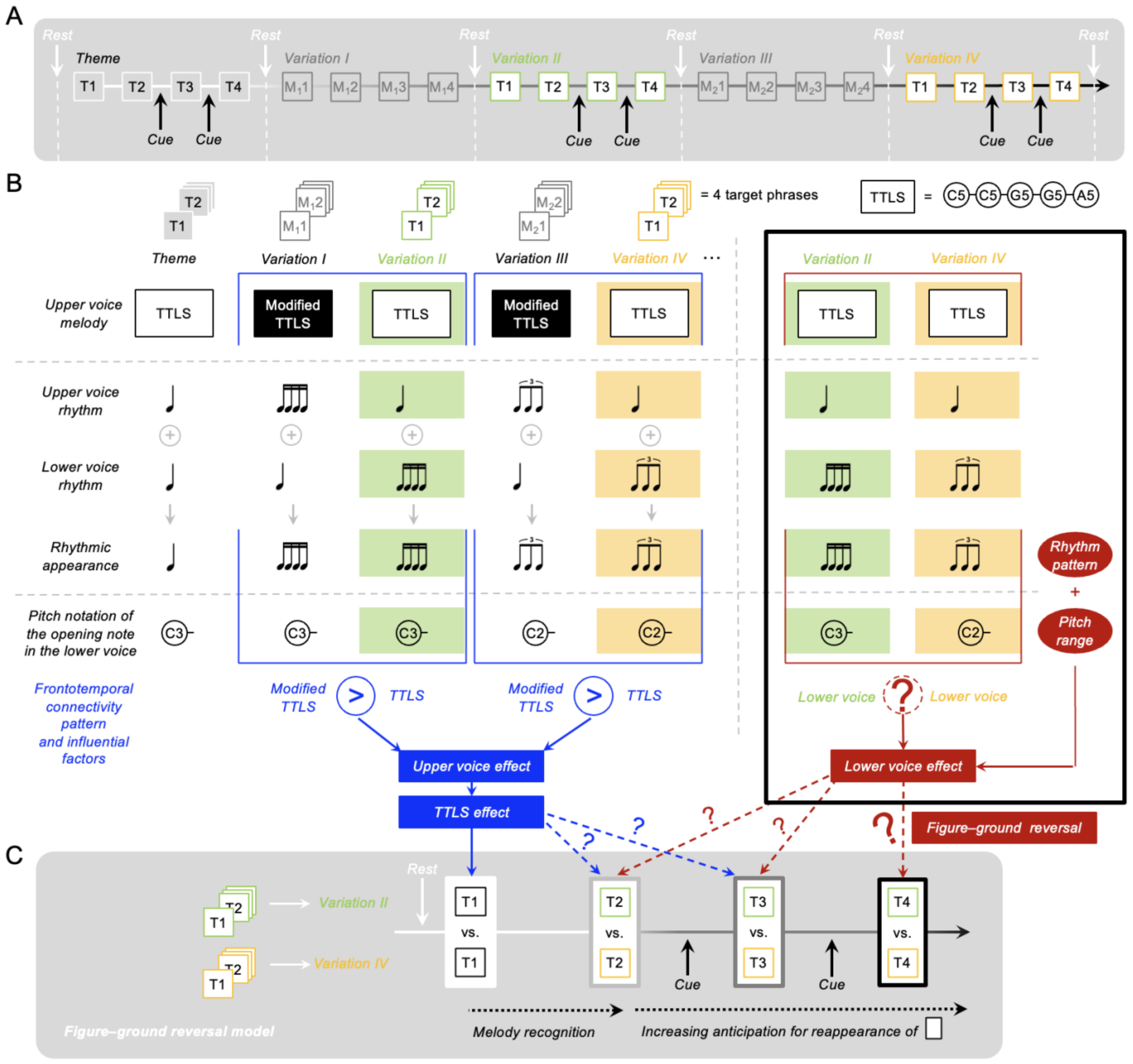


**Fig. S2. Frontotemporal connectivity and figure–ground reversal.** **(A)** Each variation involving repeat signs in the musical score for Fig. S1 is played as a ternary form, consisting of 48 measures: A (a + a) + B (b + a' + B (b + a') (Fig. 2A). A rest occurs between variations. The first modified target phrase ($M_{1}1$ and $M_{2}1$) in *Variations I* and *III*, as well as the first target phrases (T1) in the theme, *Variation II*, and *Variation IV*, are given after the rest. The phrases T1/T2, $M_{1}1$/$M_{1}2$, and $M_{2}1$/$M_{2}2$ are less predictable than T3/T4, $M_{1}3$/$M_{1}4$, and $M_{2}3$/$M_{2}4$, which are presented after the cue phrases. The TTLS melody, C5-C5-G5-G5-A5, is included in T1–T4 and modified in $M_1$ or $M_2$. **(B)** The TTLS melody in the theme, *Variation II*, and *Variation IV* is useful for observing the effects of repeated familiar melodies. *Variations I* to *IV* contain rhythmic variations on the theme. Relative to the theme, the rhythmic patterns in the upper voices for *Variations I* and *III* are transformed, while in *Variations II* and *IV*, they are moved into lower voices sharing the same TTLS melody. The rhythmic appearances and pitch ranges are consistent in each pair of *Variation I* and *II* and *Variation III* and *IV*. At T1, the frontotemporal connectivity presents a consistent pattern for both pairs, regardless of the differences in the rhythmic appearance and the range of pitches. **(C)** We defined the neuroscientific concept of figure–ground reversal as variation in frontotemporal connectivity between two variations sharing the TTLS melody but with differing lower voices, for the following reasons: (1) If the figure and ground are switched between the upper and lower voices, the role of the frontotemporal connectivity specialized in the processing upper voices, would shift to the processing of lower voices for the pairs *Variation I* vs. *II* and *Variation III* vs. *IV*. To control for the effect of the upper voice within the figure–ground reversal model, we selected four target phrases in *Variations II* and *IV* sharing the same TTLS melody. (2) Frontotemporal connectivity for *Variation II* vs. *IV* was tested in target phrases from T1 to T4. This approach allowed target phrases to be compared within the consistent context of TTLS–TTLS–cue–TTLS–cue–TTLS across both variations. 3) The difference in the frontotemporal connectivity between T1 and other Ts (i.e., T1 vs. T2, T1 vs. T3, T1 vs. T4) could indicate a change that would reflect the effect of pure repetition. Repetition is a crucial factor that influences figure–ground reversal in this study. However, as the repeated target phrases have their own context in a ternary form and are separated by a rest between variations, the target phrases should be compared in parallel between the variations. We hypothesized that, if the figure–ground relationship between voices changes between *Variations II* and *IV*, unlike the consistent frontotemporal connectivity pattern in T1, change in frontotemporal connectivity would be observed in target phrases other than T1. The frontotemporal connectivity that is evoked by the figure–ground reversal could be associated with the features of the lower voices. LTDMI strength in frontotemporal connectivity could reflect the processing of all musical elements, including rhythmic patterns and pitch ranges, which would differ in the lower voices of *Variations II* and *IV*.

**Table S1. Results of Wilcoxon signed-rank test between *Variations II* and *IV* in four target phrases for 12 connections between the bilateral HGs and IFGs.** We tested for 12 connections between the bilateral HGs and IFGs to confirm whether our hypothesis was valid, even though our hypothesis was limited to one connection among the 12 connections. Significance was observed in a single connection from the left IFG to the right HG in T4 among 48 combinations of 12 connections and four target phrases (N = 25). A tendency toward significance was observed in a connection from the right HG to the left IFG. *, $P < 0.05$.

| | **T1** | | **T2** | | **T3** | | **T4** | |
|---|---|---|---|---|---|---|---|---|
| | **Z** | **P** | **Z** | **P** | **Z** | **P** | **Z** | **P** |
| **Left HG → Right HG** | -0.296 | 0.767 | -0.578 | 0.563 | -0.632 | 0.527 | -0.632 | 0.527 |
| **Left HG → Left IFG** | -0.498 | 0.619 | -1.278 | 0.201 | -0.874 | 0.382 | -0.901 | 0.367 |
| **Left HG → Right IFG** | -0.794 | 0.427 | -0.794 | 0.427 | -1.682 | 0.093 | -0.336 | 0.737 |
| **Right HG → Left HG** | -1.278 | 0.201 | -0.363 | 0.716 | -1.332 | 0.183 | -0.283 | 0.778 |
| **Right HG → Left IFG** | -0.296 | 0.767 | -0.148 | 0.882 | -0.256 | 0.798 | ***-1.843*** | ***0.065*** |
| **Right HG → Right IFG** | -0.659 | 0.510 | -1.574 | 0.115 | -1.063 | 0.288 | -0.363 | 0.716 |
| **Left IFG → Left HG** | -0.013 | 0.989 | -0.982 | 0.326 | -1.036 | 0.300 | -1.574 | 0.115 |
| **Left IFG → Right HG** | -0.148 | 0.882 | -0.767 | 0.443 | -0.767 | 0.443 | ***-2.112*** | ***0.035 **** |
| **Left IFG → Right IFG** | -1.682 | 0.093 | -0.928 | 0.353 | -0.229 | 0.819 | -0.578 | 0.563 |
| **Right IFG → Left HG** | -1.009 | 0.313 | -0.632 | 0.527 | -0.013 | 0.989 | -0.067 | 0.946 |
| **Right IFG → Right HG** | -0.027 | 0.979 | -1.440 | 0.150 | -0.417 | 0.677 | -1.090 | 0.276 |
| **Right IFG → Left IFG** | -1.413 | 0.158 | -0.928 | 0.353 | -1.762 | 0.078 | -0.484 | 0.628 |

**Table S2. Correlation between frontotemporal connectivity and other connectivity for the difference values of *Variation II* and *IV*.** We first computed the differences in value between *Variations II* and *IV* (*Variation IV– Variation II*) for the LTDMI values in 12 connections for the bilateral IFGs and HGs for all target phrases of T1 to T4. Next, we estimated the correlations between frontotemporal connectivity difference value (Left IFG → Right HG $_{(Variation\ IV–\ Variation\ II)}$) and other 11 values for the connectivity difference. In the Spearman correlation test result, the significant positive correlation was only observed between Left IFG → Right HG $_{(Variation\ IV–\ Variation\ II)}$ and Right HG → Left IFG $_{(Variation\ IV–\ Variation\ II)}$ for T4. Type I errors caused by multiple comparisons between the 11 connection pairs for each target phrase in the Spearman correlation test were adjusted with the Bonferroni test.

| | **A** | **B** | **Correlation between A and B** | | |
|---|---|---|---|---|---|
| **Target phrase** | **Difference value of "*Variation IV – Variation II*"** | **Difference value of "*Variation IV – Variation II*"** | **Spearman's rho** | ***P*** | **Bonferroni-corrected *P*** |
| ***T1*** | Left IFG → Right HG | Right HG → Left IFG | 0.471 | 0.018 | 0.193 |
| | | Left HG → Right STG | 0.000 | 0.999 | 10.984 |
| | | Left HG → Left IFG | 0.147 | 0.483 | 5.318 |
| | | Left HG → Right IFG | -0.200 | 0.338 | 3.716 |
| | | Right HG → Left HG | 0.157 | 0.453 | 4.978 |
| | | Right HG → Right IFG | -0.113 | 0.590 | 6.495 |
| | | Left IFG → Left HG | 0.142 | 0.500 | 5.497 |
| | | Left IFG → Right IFG | 0.160 | 0.444 | 4.880 |
| | | Right IFG → Left HG | -0.267 | 0.197 | 2.168 |
| | | Right IFG → Right HG | 0.068 | 0.745 | 8.196 |
| | | Right IFG → Left IFG | 0.077 | 0.715 | 7.862 |
| ***T2*** | Left IFG → Right HG | Right HG → Left IFG | 0.515 | 0.008 | 0.092 |
| | | Left HG → Right STG | 0.293 | 0.155 | 1.706 |
| | | Left HG → Left IFG | 0.145 | 0.490 | 5.394 |
| | | Left HG → Right IFG | -0.001 | 0.996 | 10.952 |
| | | Right HG → Left HG | 0.402 | 0.046 | 0.510 |
| | | Right HG → Right IFG | -0.216 | 0.299 | 3.293 |
| | | Left IFG → Left HG | 0.005 | 0.980 | 10.776 |
| | | Left IFG → Right IFG | 0.074 | 0.724 | 7.967 |
| | | Right IFG → Left HG | -0.116 | 0.582 | 6.397 |
| | | Right IFG → Right HG | -0.052 | 0.807 | 8.874 |
| | | Right IFG → Left IFG | -0.185 | 0.375 | 4.124 |
| ***T3*** | Left IFG → Right HG | Right HG → Left IFG | 0.546 | 0.005 | 0.053 |
| | | Left HG → Right STG | 0.379 | 0.061 | 0.676 |
| | | Left HG → Left IFG | -0.457 | 0.022 | 0.239 |
| | | Left HG → Right IFG | 0.112 | 0.595 | 6.550 |
| | | Right HG → Left HG | 0.222 | 0.286 | 3.149 |
| | | Right HG → Right IFG | 0.163 | 0.437 | 4.808 |
| | | Left IFG → Left HG | -0.172 | 0.411 | 4.522 |
| | | Left IFG → Right IFG | 0.435 | 0.030 | 0.329 |
| | | Right IFG → Left HG | 0.220 | 0.290 | 3.187 |
| | | Right IFG → Right HG | 0.189 | 0.365 | 4.014 |
| | | Right IFG → Left IFG | 0.546 | 0.005 | 0.052 |
| ***T4*** | **Left IFG → Right HG** | **Right HG → Left IFG** | ***0.759*** | ***0.00001*** | ***0.0001 **** |
| | | Left HG → Right STG | 0.037 | 0.861 | 9.470 |
| | | Left HG → Left IFG | -0.255 | 0.218 | 2.396 |
| | | Left HG → Right IFG | -0.397 | 0.049 | 0.544 |

| | | | |
|---|---|---|---|
| Right HG → Left HG | 0.013 | 0.951 | 10.456 |
| Right HG → Right IFG | 0.188 | 0.369 | 4.058 |
| Left IFG → Left HG | -0.304 | 0.140 | 1.538 |
| Left IFG → Right IFG | 0.075 | 0.722 | 7.937 |
| Right IFG → Left HG | -0.022 | 0.919 | 10.105 |
| Right IFG → Right HG | 0.155 | 0.460 | 5.065 |
| Right IFG → Left IFG | 0.224 | 0.282 | 3.103 |
